# T-CoV: a comprehensive portal of HLA-peptide interactions affected by SARS-CoV-2 mutations

**DOI:** 10.1101/2021.07.06.451227

**Authors:** Stepan Nersisyan, Anton Zhiyanov, Maxim Shkurnikov, Alexander Tonevitsky

**Affiliations:** Faculty of Biology and Biotechnology, HSE University, Moscow, Russia; Shemyakin-Ovchinnikov Institute of Bioorganic Chemistry, Russian Academy of Sciences, Moscow, Russia

## Abstract

Rapidly appearing SARS-CoV-2 mutations can affect T cell epitopes, which can help the virus to evade either CD8 or CD4 T-cell responses. We developed T-cell COVID-19 Atlas (T-CoV, https://t-cov.hse.ru) – the comprehensive web portal, which allows one to analyze how SARS-CoV-2 mutations alter the presentation of viral peptides by HLA molecules. The data are presented for common virus variants and the most frequent HLA class I and class II alleles. Binding affinities of HLA molecules and viral peptides were assessed with accurate *in silico* methods. The obtained results highlight the importance of taking HLA alleles diversity into account: mutation-mediated alterations in HLA-peptide interactions were highly dependent on HLA alleles. For example, we found that the essential number of peptides tightly bound to HLA-B*07:02 in the reference Wuhan variant ceased to be tight binders for the Indian (Delta) and the UK (Alpha) variants. In summary, we believe that T-CoV will help researchers and clinicians to predict the susceptibility of individuals with different HLA genotypes to infection with variants of SARS-CoV-2 and/or forecast its severity.

## INTRODUCTION

T cells play a key role in the immune response to viral infections, including COVID-19. The development of T-cell immunity begins with a presentation of short viral peptides on a cell surface by human leukocyte antigen (HLA) complex. Within this framework, CD8 (cytotoxic, killer) T cells become activated after recognition of viral peptides presented by HLA class I molecules, which then allows them to recognize and destroy infected cells. Similarly, a receptor of CD4 T lymphocyte binds to a complex formed by a viral peptide and HLA class II molecule, which results in activation of the T-cell (1). The main function of activated CD4 (helper) T cells consists in regulation of immunity, including stimulation of antibody generation by B cells and enhancement of CD8 T-cell response (2). The important role of T-cell responses in COVID-19 severity and long-term immunity was shown in multiple reports (3–9).

Novel mutations in SARS-CoV-2 genome consistently appear and can target T-cell epitopes (10, 11). While it is hard to track the changes in interaction of T-cell receptor (TCR) and mutated epitope, the binding activity between HLA molecules and peptides could be predicted effectively *in silico* (12, 13). The main challenge here consists in the diversity of HLA alleles. Specifically, there exists dozens of common variants for each allele of HLA receptors, and each variant has its own repertoire of peptides which could be effectively presented. The distribution of HLA alleles is country and population specific (14).

In this paper we introduce T-cell COVID-19 Atlas (T-CoV) – the portal, which allows one to track the influence of SARS-CoV-2 mutations on HLA-peptide interactions. The website (https://t-cov.hse.ru) contains comprehensive data generated by bioinformatics analysis for the most prevalent SARS-CoV-2 variants and CD8/CD4 epitopes. Here we describe the main features of the portal and illustrate the possible applications for several common virus variants.

## MATERIALS AND METHODS

### SARS-CoV-2 protein and peptide data processing

Proteomes of the common SARS-CoV-2 strains (27 viral proteins) were obtained from the GISAID portal (15). As the reference virus we took a commonly used Wuhan-Hu-1 variant (GISAID accession EPI_ISL_402125). The other variants processed by the date of manuscript submission (July 2021) are listed in Supplemental Table S1.

The set of viral peptides was generated by considering all possible 8- to 14-mers from viral proteins for HLA class I receptors (potential CD8 epitopes) and 15- to 20-mers for HLA class II (CD4 epitopes). To account for mutations associated with a particular strain, we processed viral protein sequences by the following procedure. First, we constructed pairwise global alignment of the reference and mutated proteins using Biopython (16). This allowed us to build a correspondence between reference and mutated peptides by taking peptide pairs with the same start/end coordinates on the alignment (gaps were removed from peptide sequences). If reference and mutated peptides were the same, we excluded this pair from the downstream analysis. Two special cases were also analyzed:

1. Either a reference or the mutated peptide was too short to be considered as a CD8/CD4 epitope. For example, a deletion inside a reference 8-mer peptide will result in the 7-mer which is not suitable for presentation on HLA-I. In this case, the reference peptide was marked as “unpaired” (this was denoted as a gap symbol on the portal).
2. The mutated peptide belonged to the set of reference peptides (after exclusion of gaps). As an example, consider the FKLKDCVMY peptide from the NSP6 protein of the reference proteome. SGF 106-108 deletion in the UK strain resulted in the deletion of the first amino acid of the peptide, so the considered peptide takes the form KLKDCVMY after mutation. Since this sequence belonged to the reference peptidome, we considered the FKLKDCVMY peptide as “unpaired”.

### HLA alleles data

The following HLA genes were selected for the analysis: HLA-A, -B, -C (HLA class I) and HLA-DPA1/DPB1, -DQA1/DQB1, -DRB1 (HLA class II). Aiming to compose a list of the most frequent HLA alleles, we first used CIWD v3.0.0 database (17). The second field high-resolution HLA allele frequencies from different population groups such as AFA (African/African American), API (Asian/Pacific Islands), EURO (European/European descent), MENA (Middle East/North Coast of Africa), HIS (South or Central America/Hispanic/Latino), NAM (Native American populations) and UNK (unknown/not asked/multiple ancestries/other) were obtained for all mentioned genes except HLA-DPA1 and HLA-DQA1 (data were not available). Then, the union of the 10 most common alleles for each population group was taken to compose the final alleles list. The set of the most frequent HLA-DPA1 and HLA-DQA1 alleles was obtained from the PyPop database (18). The final list of considered alleles is presented in Supplemental Table S2.

### Prediction of HLA-peptide binding affinities

To predict binding affinities between the selected peptides and different HLA class I/II molecules, we used netMHCpan-4.1 and netMHCIIpan-4.0 tools, respectively (12). The predicted affinities were classified into three groups: tight binding (IC_50_ affinity ≤ 50 nM), moderate binding (50 nM < IC_50_ affinity ≤ 500 nM), weak/no binding (IC_50_ affinity > 500 nM). Note that if a reference peptide was marked as “unpaired”, we used the weak/no binding category for the mutated entry.

### Comparison of reference and mutated immunopeptidomes

For each mutated region (continuous fragment of replacements, deletions or insertions) we considered all reference/mutated peptide pairs and their interactions with frequent HLA molecules. These interactions were sorted according to affinity fold changes (i.e., differences between HLA-peptide affinity for a mutated and the reference peptides). We discarded the interactions whose affinity was not altered by at least two folds during the mutation. Finally, for each HLA allele we calculated the number of peptides with increased and decreased affinity (two or more times). Since these numbers did not account for the initial number of tightly binding peptides, we normalized the absolute numbers of new/vanished tight binders (IC_50_ affinity ≤ 50 nM) by the total number of tight binders for each allele. To avoid possible divisions by zero, we used + 1 regularization term in all denominators. For example, if some allele had no tight binders in the reference genome, and three new tight binders appeared as a result of the mutation, the relative increase will be 300%. The absolute and normalized values were presented on protein-specific and protein-aggregated plots.

### Implementation

Flask Python framework was used to build the website. Data processing and visualization were conducted with the extensive use of Pandas (19), NumPy (20), SciPy (21) and Seaborn (22) Python packages. All source codes have been made available on GitHub (https://github.com/s-a-nersisyan/TCellCovid19Atlas).

## RESULTS

### Overview of the portal

The main page of the portal (https://t-cov.hse.ru/) contains the list of the main SARS-CoV-2 strains; by the date of manuscript submission (July 2021) these were Alpha (B.1.1.7), Beta (B.1.351), Gamma (P.1), Delta (B.1.617.2+AY.1+AY.2), Epsilon (B.1.429+B.1.427), Zeta (P.2), Eta (B.1.525), Theta (P.3), Iota (B.1.526), Kappa (B.1.617.1) and Lambda (C.37) variants. As new variants possibly emerge in the future, we will promptly update the database.

For each strain we analyze CD8 and CD4 responses separately. In both cases, the top section of the page contains the set of viral mutation regions grouped by viral protein names (Figure 1A). For a single mutation (either amino acid replacement, deletion or insertion) we included the fragment of the protein sequence alignment (the reference strain against the variant of interest) and a table with HLA-peptide interactions which were significantly affected by the mutation (Figure 1B, see details in Materials and Methods).

**Figure 1.**
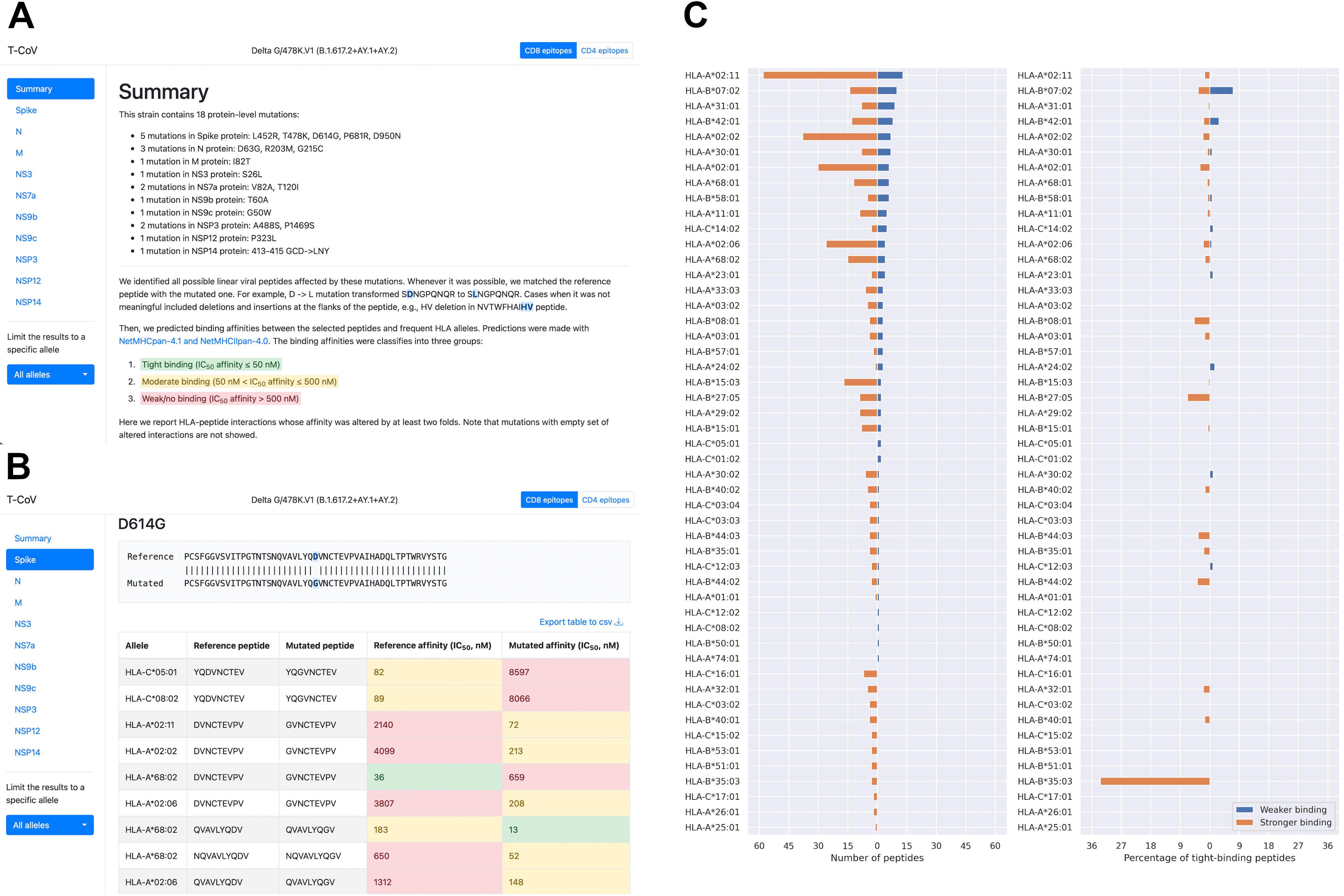
The web interface of T-CoV. **(A)** Top part of the page contains SARS-CoV-2 variant name, list of protein-level mutations, short introduction and two navigation panels: through viral proteins and different HLA alleles. **(B)** A single mutation analysis includes a fragment of pairwise sequence alignment (the reference variant and the variant of consideration) and a table with HLA-peptide interactions significantly affected by the analyzed mutation. **(C)** Allele-specific differences between numbers of T-cell epitopes from the reference virus and the variant of consideration (plot was constructed for the Delta variant). Left panel stands for the absolute number of peptides, while the right panel represents percentage of tight HLA-peptide interactions (absolute number relative to the number of tight-binders in the reference immunopeptidome).

Besides mutation-wise analysis, we present summary figures both on protein and whole virus levels. The first plot represents the numbers of HLA-peptide interactions with increased or decreased affinities (at least two folds change, Figure 1C). While absolute numbers are of interest, they do not reflect the initial number of tightly binding peptides for the specific allele. Roughly speaking, if an allele had a high number of tight binders, then vanished affinity of several peptides will not affect the integral ability of peptide presentation. To the contrary, a low number of vanished peptides could be important if the allele initially had a narrow epitope repertoire. To account for this effect, we added the second figure reflecting the ratio of the absolute numbers of new/vanished tight binders (IC50 affinity ≤ 50 nM) and the total number of tight binders for each allele (Figure 1C).

To make the page interface more friendly and comfortable to use, we provide an ability to navigate through viral peptides and restrict the analysis results to a specific HLA allele (Figure 1A). Note that all tables and plots can be exported straightforwardly from the website.

### HLA-B*07:02 is a potential risk allele for the Indian and the UK strains

We previously mentioned that the absolute number of peptides with stronger or weaker binding affinities with a specific allele may not reflect the real biological effects, since the latter strongly depends on the total number of epitopes in the reference virus. To illustrate this idea, we refer to the Indian Delta variant (B.1.617.2+AY.1+AY.2). Considering the allele HLA-A*02:11, we note that the absolute number of peptides with altered binding affinity was very high (13 peptides with weaker and 58 peptides with stronger binding), while the initial number of tight binding peptides was 1072: only 1 of them became non-tight binders (0.1%) and 17 novel tight binders appeared (1.6%). Thus, the mutations of the Indian strain will probably not affect much the peptide presentation ability of HLA-A*02:11 despite the high absolute number of predicted changes.

In this context we highlight HLA-B*07:02 allele. Because of the P681R mutation in the Spike protein, affinity of four tightly binding peptides sharply decreased (7.1% of the total tight binders). Interestingly, the similar situation was observed in the context of the UK Alpha variant (B.1.1.7). Namely, four tight HLA-peptide interactions vanished, however, this was caused by the different mutation (P681H, Spike protein). Note, that HLA-B*07:02 allele is one of the most frequent alleles of HLA-B in the world, especially in the Sweden (19%), Ireland (17%), UK (15%), Netherlands (15%), Russia (7-13%), Austria (13%) and Germany (12%) according to The Allele Frequency Net Database (23).

Aside from the aforementioned examples, some cases were associated with significant changes both in terms of absolute numbers and percentages of tight binders. Specifically, mutations in the Spike protein (the UK variant) led to the weaker binding of 81 peptides and HLA-DRB1*15:01, including loss of 22 tight binders (15%). Remarkably, all vanished tight binders were affected by the HV 69-70 deletion.

### Premature ORF8 stop codon in the UK strain results in loss of multiple epitopes

Among all mutations of the UK strain, the appearance of the premature stop codon in the ORF8 (NS8) protein (Q27stop) is of particular interest since it results in the deletion of almost 80% of the protein. Even though functional consequences of this mutation are not fully understood (24), we note that the whole immunopeptidome associated with this part of protein entirely vanished. In the context of CD8 epitopes, this included 46 strong (19.8%) and 186 moderate (80.2%) interactions of HLA-I molecules and peptides. The similar situation was observed for HLA class II and CD4 epitopes: 315 pairs with tight binding activity (4.1%), 7435 moderate pairs (95.9%) became weaker binders.

## DISCUSSION

We developed the web portal for the analysis of changes in HLA-peptide binding caused by mutations of SARS-CoV-2. Comprehensive analysis was performed for the major SARS-CoV-2 strains and the most frequent HLA class I/II alleles. The influence of mutations on viral peptide presentation by HLA molecules was very allele-specific, indicating the potential existence of individuals which could be susceptible to severe COVID-19 caused by a particular virus strain.

There is an experimental evidence on the negative role of virus mutations on T-cell immunity. In a recent work Reynolds with co-authors studied CD4 T cell immune response to the UK and the South African variants after first dose vaccinations among health care workers with or without prior Wuhan-Hu-1 SARS-CoV-2 infection (11). It turned out that T cell responses to the variant peptides associated with D1118H mutation of Spike protein were reduced in the individuals carrying HLA-DRB1*03:01 allele. There is a basis for this in terms of differential HLA class II binding: all HLA-peptide pairs associated with this mutation moved from the moderate binding category to the weak/no binding group. In another report, Agerer et al found reduced binding of mutated peptides to HLA-A*02:01 and HLA-B*40:01 alleles, the most frequent mutations were inferred from the Austrian patients cohort (10). As suggested by our analysis, mutation-mediated effects are HLA allele specific. Thus, more experiments are required to cover the essential part of the diverse HLA molecules set.

We note several limitations of our work. The main limitation is since successful peptide presentation by HLA molecule does not guarantee further recognition by a TCR. Thus, clinical implications of significant alterations in HLA-peptide binding affinity should be further experimentally assessed. The second limitation involves officially unannotated SARS-CoV-2 proteins which could contain T cell epitopes: proteomics analysis carried out by Weingarten-Gabbay with co-authors resulted in detection of highly affine HLA-I peptides from out-of-frame ORFs (25). While some of these ORFs were included in our analysis (e.g., ORF9b), the other entries like S.iORF1/2 were not considered. Moreover, synonymous nucleotide substitutions which were ignored in our study could result in nonsynonymous amino acid substitutions in noncanonical reading frames. Such kind of analysis will be included in the next versions of T-CoV portal.

## AVAILABILITY

T-CoV portal is available at the following link: https://t-cov.hse.ru. All source codes have been made available on GitHub (https://github.com/s-a-nersisyan/TCellCovid19Atlas).

## ACKNOWLEDGEMENT

The authors thank Milena Chekova and Narek Engibaryan for the help with composition of the list of most frequent HLA alleles.

