## Supplementary Data for "T-CoV: a comprehensive portal of HLA-peptide interactions affected by SARS-CoV-2 mutations"

**Supplemental Table S1. SARS-CoV-2 variants used in the analysis by the date of manuscript submission (July 2021).**

| **Variant** | **GISAID virus name** | **GISAID accession ID** |
| --- | --- | --- |
| Alpha 202012/01 GRY (B.1.1.7) | hCoV-19/England/MILK-9E2FE0/2020 | EPI_ISL_581117 |
| Beta GH/501Y.V2 (B.1.351) | hCoV-19/South Africa/KRISP-K004312/2020 | EPI_ISL_660190 |
| Gamma GR/501Y.V3 (P.1) | hCoV-19/Japan/IC-0561/2021 | EPI_ISL_792680 |
| Delta G/478K.V1 (B.1.617.2+AY.1+AY.2) | hCoV-19/India/ILSGS00941/2020 | EPI_ISL_1663516 |
| Epsilon GH/452R.V1 (B.1.429+B.1.427) | hCoV-19/USA/CA-CZB-12872/2020 | EPI_ISL_648527 |
| Zeta GR/484K.V2 (P.2) | hCoV-19/England/PORT-2EAFC1/2020 | EPI_ISL_842052 |
| Eta G/484K.V3 (B.1.525) | hCoV-19/England/CAMC-C769B3/2020 | EPI_ISL_760883 |
| Theta GR/1092K.V1 (P.3) | hCoV-19/Hong Kong/CM21000064/2021 | EPI_ISL_1914574 |
| Iota GH/253G.V1 (B.1.526) | hCoV-19/USA/NY-MSHSPSP-PV21166/2020 | EPI_ISL_801973 |
| Kappa G/452R.V3 (B.1.617.1) | hCoV-19/India/ILSGS00308/2020 | EPI_ISL_1372093 |
| Lambda GR/452Q.V1 (C.37) | hCoV-19/Peru/LIM-INS-869/2020 | EPI_ISL_1534645 |

**Supplemental Table S2. HLA alleles used in the study.**

| **HLA allele** | **Gene** | **HLA class** |
| --- | --- | --- |
| HLA-A*01:01 | HLA-A | I |
| HLA-A*02:01 | HLA-A | I |
| HLA-A*02:02 | HLA-A | I |
| HLA-A*02:06 | HLA-A | I |
| HLA-A*02:11 | HLA-A | I |
| HLA-A*03:01 | HLA-A | I |
| HLA-A*03:02 | HLA-A | I |
| HLA-A*11:01 | HLA-A | I |
| HLA-A*23:01 | HLA-A | I |
| HLA-A*24:02 | HLA-A | I |
| HLA-A*25:01 | HLA-A | I |
| HLA-A*26:01 | HLA-A | I |
| HLA-A*29:02 | HLA-A | I |
| HLA-A*30:01 | HLA-A | I |
| HLA-A*30:02 | HLA-A | I |
| HLA-A*31:01 | HLA-A | I |
| HLA-A*32:01 | HLA-A | I |
| HLA-A*33:03 | HLA-A | I |
| HLA-A*68:01 | HLA-A | I |
| HLA-A*68:02 | HLA-A | I |
| HLA-A*74:01 | HLA-A | I |
| HLA-B*07:02 | HLA-B | I |
| HLA-B*08:01 | HLA-B | I |
| HLA-B*14:02 | HLA-B | I |
| HLA-B*15:01 | HLA-B | I |
| HLA-B*15:03 | HLA-B | I |
| HLA-B*18:01 | HLA-B | I |
| HLA-B*27:05 | HLA-B | I |
| HLA-B*35:01 | HLA-B | I |
| HLA-B*35:03 | HLA-B | I |
| HLA-B*38:01 | HLA-B | I |
| HLA-B*40:01 | HLA-B | I |
| HLA-B*40:02 | HLA-B | I |
| HLA-B*40:06 | HLA-B | I |
| HLA-B*42:01 | HLA-B | I |
| HLA-B*44:02 | HLA-B | I |
| HLA-B*44:03 | HLA-B | I |
| HLA-B*45:01 | HLA-B | I |
| HLA-B*50:01 | HLA-B | I |
| HLA-B*51:01 | HLA-B | I |
| HLA-B*52:01 | HLA-B | I |
| HLA-B*53:01 | HLA-B | I |
| HLA-B*57:01 | HLA-B | I |
| HLA-B*58:01 | HLA-B | I |
| HLA-B*58:02 | HLA-B | I |
| HLA-C*01:02 | HLA-C | I |
| HLA-C*02:02 | HLA-C | I |
| HLA-C*02:10 | HLA-C | I |
| HLA-C*03:02 | HLA-C | I |
| HLA-C*03:03 | HLA-C | I |
| HLA-C*03:04 | HLA-C | I |
| HLA-C*04:01 | HLA-C | I |
| HLA-C*05:01 | HLA-C | I |
| HLA-C*06:02 | HLA-C | I |
| HLA-C*07:01 | HLA-C | I |
| HLA-C*07:02 | HLA-C | I |
| HLA-C*08:01 | HLA-C | I |
| HLA-C*08:02 | HLA-C | I |
| HLA-C*12:02 | HLA-C | I |
| HLA-C*12:03 | HLA-C | I |
| HLA-C*14:02 | HLA-C | I |
| HLA-C*15:02 | HLA-C | I |
| HLA-C*16:01 | HLA-C | I |
| HLA-C*17:01 | HLA-C | I |
| HLA-DRB1*01:01 | HLA-DRB1 | II |
| HLA-DRB1*01:02 | HLA-DRB1 | II |
| HLA-DRB1*03:01 | HLA-DRB1 | II |
| HLA-DRB1*03:02 | HLA-DRB1 | II |
| HLA-DRB1*04:01 | HLA-DRB1 | II |
| HLA-DRB1*04:02 | HLA-DRB1 | II |
| HLA-DRB1*04:03 | HLA-DRB1 | II |
| HLA-DRB1*04:04 | HLA-DRB1 | II |
| HLA-DRB1*04:07 | HLA-DRB1 | II |
| HLA-DRB1*07:01 | HLA-DRB1 | II |
| HLA-DRB1*08:02 | HLA-DRB1 | II |
| HLA-DRB1*08:04 | HLA-DRB1 | II |
| HLA-DRB1*10:01 | HLA-DRB1 | II |
| HLA-DRB1*11:01 | HLA-DRB1 | II |
| HLA-DRB1*11:02 | HLA-DRB1 | II |
| HLA-DRB1*11:04 | HLA-DRB1 | II |
| HLA-DRB1*12:02 | HLA-DRB1 | II |
| HLA-DRB1*13:01 | HLA-DRB1 | II |
| HLA-DRB1*13:02 | HLA-DRB1 | II |
| HLA-DRB1*14:01 | HLA-DRB1 | II |
| HLA-DRB1*14:04 | HLA-DRB1 | II |
| HLA-DRB1*15:01 | HLA-DRB1 | II |
| HLA-DRB1*15:02 | HLA-DRB1 | II |
| HLA-DRB1*15:03 | HLA-DRB1 | II |
| HLA-DRB1*16:01 | HLA-DRB1 | II |
| HLA-DPA1*01:03/DPB1*01:01 | HLA-DPA1/DPB1 | II |
| HLA-DPA1*02:01/DPB1*01:01 | HLA-DPA1/DPB1 | II |
| HLA-DPA1*02:02/DPB1*01:01 | HLA-DPA1/DPB1 | II |
| HLA-DPA1*03:01/DPB1*01:01 | HLA-DPA1/DPB1 | II |
| HLA-DPA1*04:01/DPB1*01:01 | HLA-DPA1/DPB1 | II |
| HLA-DPA1*01:03/DPB1*02:01 | HLA-DPA1/DPB1 | II |
| HLA-DPA1*02:01/DPB1*02:01 | HLA-DPA1/DPB1 | II |
| HLA-DPA1*02:02/DPB1*02:01 | HLA-DPA1/DPB1 | II |
| HLA-DPA1*03:01/DPB1*02:01 | HLA-DPA1/DPB1 | II |
| HLA-DPA1*04:01/DPB1*02:01 | HLA-DPA1/DPB1 | II |
| HLA-DPA1*01:03/DPB1*03:01 | HLA-DPA1/DPB1 | II |
| HLA-DPA1*02:01/DPB1*03:01 | HLA-DPA1/DPB1 | II |
| HLA-DPA1*02:02/DPB1*03:01 | HLA-DPA1/DPB1 | II |
| HLA-DPA1*03:01/DPB1*03:01 | HLA-DPA1/DPB1 | II |
| HLA-DPA1*04:01/DPB1*03:01 | HLA-DPA1/DPB1 | II |
| HLA-DPA1*01:03/DPB1*04:01 | HLA-DPA1/DPB1 | II |
| HLA-DPA1*02:01/DPB1*04:01 | HLA-DPA1/DPB1 | II |
| HLA-DPA1*02:02/DPB1*04:01 | HLA-DPA1/DPB1 | II |
| HLA-DPA1*03:01/DPB1*04:01 | HLA-DPA1/DPB1 | II |
| HLA-DPA1*04:01/DPB1*04:01 | HLA-DPA1/DPB1 | II |
| HLA-DPA1*01:03/DPB1*04:02 | HLA-DPA1/DPB1 | II |
| HLA-DPA1*02:01/DPB1*04:02 | HLA-DPA1/DPB1 | II |
| HLA-DPA1*02:02/DPB1*04:02 | HLA-DPA1/DPB1 | II |
| HLA-DPA1*03:01/DPB1*04:02 | HLA-DPA1/DPB1 | II |
| HLA-DPA1*04:01/DPB1*04:02 | HLA-DPA1/DPB1 | II |
| HLA-DPA1*01:03/DPB1*05:01 | HLA-DPA1/DPB1 | II |
| HLA-DPA1*02:01/DPB1*05:01 | HLA-DPA1/DPB1 | II |
| HLA-DPA1*02:02/DPB1*05:01 | HLA-DPA1/DPB1 | II |
| HLA-DPA1*03:01/DPB1*05:01 | HLA-DPA1/DPB1 | II |
| HLA-DPA1*04:01/DPB1*05:01 | HLA-DPA1/DPB1 | II |
| HLA-DPA1*01:03/DPB1*06:01 | HLA-DPA1/DPB1 | II |
| HLA-DPA1*02:01/DPB1*06:01 | HLA-DPA1/DPB1 | II |
| HLA-DPA1*02:02/DPB1*06:01 | HLA-DPA1/DPB1 | II |
| HLA-DPA1*03:01/DPB1*06:01 | HLA-DPA1/DPB1 | II |
| HLA-DPA1*04:01/DPB1*06:01 | HLA-DPA1/DPB1 | II |
| HLA-DPA1*01:03/DPB1*09:01 | HLA-DPA1/DPB1 | II |
| HLA-DPA1*02:01/DPB1*09:01 | HLA-DPA1/DPB1 | II |
| HLA-DPA1*02:02/DPB1*09:01 | HLA-DPA1/DPB1 | II |
| HLA-DPA1*03:01/DPB1*09:01 | HLA-DPA1/DPB1 | II |
| HLA-DPA1*04:01/DPB1*09:01 | HLA-DPA1/DPB1 | II |
| HLA-DPA1*01:03/DPB1*10:01 | HLA-DPA1/DPB1 | II |
| HLA-DPA1*02:01/DPB1*10:01 | HLA-DPA1/DPB1 | II |
| HLA-DPA1*02:02/DPB1*10:01 | HLA-DPA1/DPB1 | II |
| HLA-DPA1*03:01/DPB1*10:01 | HLA-DPA1/DPB1 | II |
| HLA-DPA1*04:01/DPB1*10:01 | HLA-DPA1/DPB1 | II |
| HLA-DPA1*01:03/DPB1*105:01 | HLA-DPA1/DPB1 | II |
| HLA-DPA1*02:01/DPB1*105:01 | HLA-DPA1/DPB1 | II |
| HLA-DPA1*02:02/DPB1*105:01 | HLA-DPA1/DPB1 | II |
| HLA-DPA1*03:01/DPB1*105:01 | HLA-DPA1/DPB1 | II |
| HLA-DPA1*04:01/DPB1*105:01 | HLA-DPA1/DPB1 | II |
| HLA-DPA1*01:03/DPB1*11:01 | HLA-DPA1/DPB1 | II |
| HLA-DPA1*02:01/DPB1*11:01 | HLA-DPA1/DPB1 | II |
| HLA-DPA1*02:02/DPB1*11:01 | HLA-DPA1/DPB1 | II |
| HLA-DPA1*03:01/DPB1*11:01 | HLA-DPA1/DPB1 | II |
| HLA-DPA1*04:01/DPB1*11:01 | HLA-DPA1/DPB1 | II |
| HLA-DPA1*01:03/DPB1*13:01 | HLA-DPA1/DPB1 | II |
| HLA-DPA1*02:01/DPB1*13:01 | HLA-DPA1/DPB1 | II |
| HLA-DPA1*02:02/DPB1*13:01 | HLA-DPA1/DPB1 | II |
| HLA-DPA1*03:01/DPB1*13:01 | HLA-DPA1/DPB1 | II |
| HLA-DPA1*04:01/DPB1*13:01 | HLA-DPA1/DPB1 | II |
| HLA-DPA1*01:03/DPB1*14:01 | HLA-DPA1/DPB1 | II |
| HLA-DPA1*02:01/DPB1*14:01 | HLA-DPA1/DPB1 | II |
| HLA-DPA1*02:02/DPB1*14:01 | HLA-DPA1/DPB1 | II |
| HLA-DPA1*03:01/DPB1*14:01 | HLA-DPA1/DPB1 | II |
| HLA-DPA1*04:01/DPB1*14:01 | HLA-DPA1/DPB1 | II |
| HLA-DPA1*01:03/DPB1*17:01 | HLA-DPA1/DPB1 | II |
| HLA-DPA1*02:01/DPB1*17:01 | HLA-DPA1/DPB1 | II |
| HLA-DPA1*02:02/DPB1*17:01 | HLA-DPA1/DPB1 | II |
| HLA-DPA1*03:01/DPB1*17:01 | HLA-DPA1/DPB1 | II |
| HLA-DPA1*04:01/DPB1*17:01 | HLA-DPA1/DPB1 | II |
| HLA-DPA1*01:03/DPB1*26:01 | HLA-DPA1/DPB1 | II |
| HLA-DPA1*02:01/DPB1*26:01 | HLA-DPA1/DPB1 | II |
| HLA-DPA1*02:02/DPB1*26:01 | HLA-DPA1/DPB1 | II |
| HLA-DPA1*03:01/DPB1*26:01 | HLA-DPA1/DPB1 | II |
| HLA-DPA1*04:01/DPB1*26:01 | HLA-DPA1/DPB1 | II |
| HLA-DPA1*01:03/DPB1*85:01 | HLA-DPA1/DPB1 | II |
| HLA-DPA1*02:01/DPB1*85:01 | HLA-DPA1/DPB1 | II |
| HLA-DPA1*02:02/DPB1*85:01 | HLA-DPA1/DPB1 | II |
| HLA-DPA1*03:01/DPB1*85:01 | HLA-DPA1/DPB1 | II |
| HLA-DPA1*04:01/DPB1*85:01 | HLA-DPA1/DPB1 | II |
| HLA-DQA1*01:01/DQB1*02:01 | HLA-DQA1/DQB1 | II |
| HLA-DQA1*01:02/DQB1*02:01 | HLA-DQA1/DQB1 | II |
| HLA-DQA1*01:03/DQB1*02:01 | HLA-DQA1/DQB1 | II |
| HLA-DQA1*02:01/DQB1*02:01 | HLA-DQA1/DQB1 | II |
| HLA-DQA1*03:01/DQB1*02:01 | HLA-DQA1/DQB1 | II |
| HLA-DQA1*04:01/DQB1*02:01 | HLA-DQA1/DQB1 | II |
| HLA-DQA1*05:01/DQB1*02:01 | HLA-DQA1/DQB1 | II |
| HLA-DQA1*06:01/DQB1*02:01 | HLA-DQA1/DQB1 | II |
| HLA-DQA1*01:01/DQB1*02:02 | HLA-DQA1/DQB1 | II |
| HLA-DQA1*01:02/DQB1*02:02 | HLA-DQA1/DQB1 | II |
| HLA-DQA1*01:03/DQB1*02:02 | HLA-DQA1/DQB1 | II |
| HLA-DQA1*02:01/DQB1*02:02 | HLA-DQA1/DQB1 | II |
| HLA-DQA1*03:01/DQB1*02:02 | HLA-DQA1/DQB1 | II |
| HLA-DQA1*04:01/DQB1*02:02 | HLA-DQA1/DQB1 | II |
| HLA-DQA1*05:01/DQB1*02:02 | HLA-DQA1/DQB1 | II |
| HLA-DQA1*06:01/DQB1*02:02 | HLA-DQA1/DQB1 | II |
| HLA-DQA1*01:01/DQB1*03:01 | HLA-DQA1/DQB1 | II |
| HLA-DQA1*01:02/DQB1*03:01 | HLA-DQA1/DQB1 | II |
| HLA-DQA1*01:03/DQB1*03:01 | HLA-DQA1/DQB1 | II |
| HLA-DQA1*02:01/DQB1*03:01 | HLA-DQA1/DQB1 | II |
| HLA-DQA1*03:01/DQB1*03:01 | HLA-DQA1/DQB1 | II |
| HLA-DQA1*04:01/DQB1*03:01 | HLA-DQA1/DQB1 | II |
| HLA-DQA1*05:01/DQB1*03:01 | HLA-DQA1/DQB1 | II |
| HLA-DQA1*06:01/DQB1*03:01 | HLA-DQA1/DQB1 | II |
| HLA-DQA1*01:01/DQB1*03:02 | HLA-DQA1/DQB1 | II |
| HLA-DQA1*01:02/DQB1*03:02 | HLA-DQA1/DQB1 | II |
| HLA-DQA1*01:03/DQB1*03:02 | HLA-DQA1/DQB1 | II |
| HLA-DQA1*02:01/DQB1*03:02 | HLA-DQA1/DQB1 | II |
| HLA-DQA1*03:01/DQB1*03:02 | HLA-DQA1/DQB1 | II |
| HLA-DQA1*04:01/DQB1*03:02 | HLA-DQA1/DQB1 | II |
| HLA-DQA1*05:01/DQB1*03:02 | HLA-DQA1/DQB1 | II |
| HLA-DQA1*06:01/DQB1*03:02 | HLA-DQA1/DQB1 | II |
| HLA-DQA1*01:01/DQB1*03:03 | HLA-DQA1/DQB1 | II |
| HLA-DQA1*01:02/DQB1*03:03 | HLA-DQA1/DQB1 | II |
| HLA-DQA1*01:03/DQB1*03:03 | HLA-DQA1/DQB1 | II |
| HLA-DQA1*02:01/DQB1*03:03 | HLA-DQA1/DQB1 | II |
| HLA-DQA1*03:01/DQB1*03:03 | HLA-DQA1/DQB1 | II |
| HLA-DQA1*04:01/DQB1*03:03 | HLA-DQA1/DQB1 | II |
| HLA-DQA1*05:01/DQB1*03:03 | HLA-DQA1/DQB1 | II |
| HLA-DQA1*06:01/DQB1*03:03 | HLA-DQA1/DQB1 | II |
| HLA-DQA1*01:01/DQB1*03:19 | HLA-DQA1/DQB1 | II |
| HLA-DQA1*01:02/DQB1*03:19 | HLA-DQA1/DQB1 | II |
| HLA-DQA1*01:03/DQB1*03:19 | HLA-DQA1/DQB1 | II |
| HLA-DQA1*02:01/DQB1*03:19 | HLA-DQA1/DQB1 | II |
| HLA-DQA1*03:01/DQB1*03:19 | HLA-DQA1/DQB1 | II |
| HLA-DQA1*04:01/DQB1*03:19 | HLA-DQA1/DQB1 | II |
| HLA-DQA1*05:01/DQB1*03:19 | HLA-DQA1/DQB1 | II |
| HLA-DQA1*06:01/DQB1*03:19 | HLA-DQA1/DQB1 | II |
| HLA-DQA1*01:01/DQB1*04:02 | HLA-DQA1/DQB1 | II |
| HLA-DQA1*01:02/DQB1*04:02 | HLA-DQA1/DQB1 | II |
| HLA-DQA1*01:03/DQB1*04:02 | HLA-DQA1/DQB1 | II |
| HLA-DQA1*02:01/DQB1*04:02 | HLA-DQA1/DQB1 | II |
| HLA-DQA1*03:01/DQB1*04:02 | HLA-DQA1/DQB1 | II |
| HLA-DQA1*04:01/DQB1*04:02 | HLA-DQA1/DQB1 | II |
| HLA-DQA1*05:01/DQB1*04:02 | HLA-DQA1/DQB1 | II |
| HLA-DQA1*06:01/DQB1*04:02 | HLA-DQA1/DQB1 | II |
| HLA-DQA1*01:01/DQB1*05:01 | HLA-DQA1/DQB1 | II |
| HLA-DQA1*01:02/DQB1*05:01 | HLA-DQA1/DQB1 | II |
| HLA-DQA1*01:03/DQB1*05:01 | HLA-DQA1/DQB1 | II |
| HLA-DQA1*02:01/DQB1*05:01 | HLA-DQA1/DQB1 | II |
| HLA-DQA1*03:01/DQB1*05:01 | HLA-DQA1/DQB1 | II |
| HLA-DQA1*04:01/DQB1*05:01 | HLA-DQA1/DQB1 | II |
| HLA-DQA1*05:01/DQB1*05:01 | HLA-DQA1/DQB1 | II |
| HLA-DQA1*06:01/DQB1*05:01 | HLA-DQA1/DQB1 | II |
| HLA-DQA1*01:01/DQB1*05:02 | HLA-DQA1/DQB1 | II |
| HLA-DQA1*01:02/DQB1*05:02 | HLA-DQA1/DQB1 | II |
| HLA-DQA1*01:03/DQB1*05:02 | HLA-DQA1/DQB1 | II |
| HLA-DQA1*02:01/DQB1*05:02 | HLA-DQA1/DQB1 | II |
| HLA-DQA1*03:01/DQB1*05:02 | HLA-DQA1/DQB1 | II |
| HLA-DQA1*04:01/DQB1*05:02 | HLA-DQA1/DQB1 | II |
| HLA-DQA1*05:01/DQB1*05:02 | HLA-DQA1/DQB1 | II |
| HLA-DQA1*06:01/DQB1*05:02 | HLA-DQA1/DQB1 | II |
| HLA-DQA1*01:01/DQB1*05:03 | HLA-DQA1/DQB1 | II |
| HLA-DQA1*01:02/DQB1*05:03 | HLA-DQA1/DQB1 | II |
| HLA-DQA1*01:03/DQB1*05:03 | HLA-DQA1/DQB1 | II |
| HLA-DQA1*02:01/DQB1*05:03 | HLA-DQA1/DQB1 | II |
| HLA-DQA1*03:01/DQB1*05:03 | HLA-DQA1/DQB1 | II |
| HLA-DQA1*04:01/DQB1*05:03 | HLA-DQA1/DQB1 | II |
| HLA-DQA1*05:01/DQB1*05:03 | HLA-DQA1/DQB1 | II |
| HLA-DQA1*06:01/DQB1*05:03 | HLA-DQA1/DQB1 | II |
| HLA-DQA1*01:01/DQB1*06:01 | HLA-DQA1/DQB1 | II |
| HLA-DQA1*01:02/DQB1*06:01 | HLA-DQA1/DQB1 | II |
| HLA-DQA1*01:03/DQB1*06:01 | HLA-DQA1/DQB1 | II |
| HLA-DQA1*02:01/DQB1*06:01 | HLA-DQA1/DQB1 | II |
| HLA-DQA1*03:01/DQB1*06:01 | HLA-DQA1/DQB1 | II |
| HLA-DQA1*04:01/DQB1*06:01 | HLA-DQA1/DQB1 | II |
| HLA-DQA1*05:01/DQB1*06:01 | HLA-DQA1/DQB1 | II |
| HLA-DQA1*06:01/DQB1*06:01 | HLA-DQA1/DQB1 | II |
| HLA-DQA1*01:01/DQB1*06:02 | HLA-DQA1/DQB1 | II |
| HLA-DQA1*01:02/DQB1*06:02 | HLA-DQA1/DQB1 | II |
| HLA-DQA1*01:03/DQB1*06:02 | HLA-DQA1/DQB1 | II |
| HLA-DQA1*02:01/DQB1*06:02 | HLA-DQA1/DQB1 | II |
| HLA-DQA1*03:01/DQB1*06:02 | HLA-DQA1/DQB1 | II |
| HLA-DQA1*04:01/DQB1*06:02 | HLA-DQA1/DQB1 | II |
| HLA-DQA1*05:01/DQB1*06:02 | HLA-DQA1/DQB1 | II |
| HLA-DQA1*06:01/DQB1*06:02 | HLA-DQA1/DQB1 | II |
| HLA-DQA1*01:01/DQB1*06:03 | HLA-DQA1/DQB1 | II |
| HLA-DQA1*01:02/DQB1*06:03 | HLA-DQA1/DQB1 | II |
| HLA-DQA1*01:03/DQB1*06:03 | HLA-DQA1/DQB1 | II |
| HLA-DQA1*02:01/DQB1*06:03 | HLA-DQA1/DQB1 | II |
| HLA-DQA1*03:01/DQB1*06:03 | HLA-DQA1/DQB1 | II |
| HLA-DQA1*04:01/DQB1*06:03 | HLA-DQA1/DQB1 | II |
| HLA-DQA1*05:01/DQB1*06:03 | HLA-DQA1/DQB1 | II |
| HLA-DQA1*06:01/DQB1*06:03 | HLA-DQA1/DQB1 | II |
| HLA-DQA1*01:01/DQB1*06:04 | HLA-DQA1/DQB1 | II |
| HLA-DQA1*01:02/DQB1*06:04 | HLA-DQA1/DQB1 | II |
| HLA-DQA1*01:03/DQB1*06:04 | HLA-DQA1/DQB1 | II |
| HLA-DQA1*02:01/DQB1*06:04 | HLA-DQA1/DQB1 | II |
| HLA-DQA1*03:01/DQB1*06:04 | HLA-DQA1/DQB1 | II |
| HLA-DQA1*04:01/DQB1*06:04 | HLA-DQA1/DQB1 | II |
| HLA-DQA1*05:01/DQB1*06:04 | HLA-DQA1/DQB1 | II |
| HLA-DQA1*06:01/DQB1*06:04 | HLA-DQA1/DQB1 | II |
| HLA-DQA1*01:01/DQB1*06:09 | HLA-DQA1/DQB1 | II |
| HLA-DQA1*01:02/DQB1*06:09 | HLA-DQA1/DQB1 | II |
| HLA-DQA1*01:03/DQB1*06:09 | HLA-DQA1/DQB1 | II |
| HLA-DQA1*02:01/DQB1*06:09 | HLA-DQA1/DQB1 | II |
| HLA-DQA1*03:01/DQB1*06:09 | HLA-DQA1/DQB1 | II |
| HLA-DQA1*04:01/DQB1*06:09 | HLA-DQA1/DQB1 | II |
| HLA-DQA1*05:01/DQB1*06:09 | HLA-DQA1/DQB1 | II |
| HLA-DQA1*06:01/DQB1*06:09 | HLA-DQA1/DQB1 | II |
